# Seascape genomics reveals limited dispersal and suggest spatially varying selection among European populations of sea lamprey (*Petromyzon marinus*)

**DOI:** 10.1101/2022.09.28.509639

**Authors:** Miguel Baltazar-Soares, J. Robert Britton, Adrian Pinder, Andrew J. Harrison, Andrew D. Nunn, Bernardo R. Quintella, Catarina S. Mateus, Jonathan D. Bolland, Jamie R. Dodd, Pedro R. Almeida, Victoria Dominguez Almela, Demetra Andreou

**Affiliations:** Department of Life and Environmental Sciences, Faculty of Science and Technology, Bournemouth University, Dorset, UK; MARE – Marine and Environmental Sciences Centre, ISPA – Instituto Universitário, Lisboa, Portugal; MARE—Marine and Environmental Sciences Centre, University of Évora, Évora, Portugal; University of Hull, Hull International Fisheries Institute, UK; Department of Animal Biology, University of Lisbon, Faculty of Sciences, Lisbon, Portugal; Department of Biology, School of Sciences and Technology, University of Évora, Évora, Portugal

**Keywords:** *Petromyzon marinus*, freshwater biodiversity, limited dispersal, seascape genomics

## Abstract

Sea lamprey *Petromyzon marinus* is an anadromous and semelparous fish without homing behaviours. Despite being a freshwater, free-living organism for large part of their life cycle, its adulthood is spent as a parasite of marine vertebrates. In their native European range, while it is well-established that sea lampreys comprise a single nearly-panmictic population, few studies have further explored the evolutionary history of natural populations. Here, we performed the first genome-wide characterization of sea lamprey’s genetic diversity in their European natural range. The objectives were to investigate the connectivity among river basins and explore evolutionary processes mediating dispersal during the marine phase, with the sequencing 186 individuals from 8 locations spanning the North Eastern Atlantic coast and the North Sea with double-digest RAD-sequencing, obtaining a total of 30910 bi-allelic SNPs. Population genetic analyses reinforced the existence of a single metapopulation encompassing freshwater spawning sites within the north eastern Atlantic and the North Sea, though the prevalence of private alleles at northern latitudes suggested some limits to the species’ dispersal. Seascape genomics suggested a scenario where oxygen concentration and river runoffs impose spatially varying selection across their distribution range. Exploring associations with abundance of potential hosts further suggested that hake and cod could also impose selective pressures, although the nature of such putative biotic interactions was unresolved. Overall, the identification of adaptive seascapes in a panmictic anadromous species could contribute to conservation practices by providing information for restoration activities to mitigate local extinctions on freshwater sites.

Sea lamprey *Petromyzon marinus* is an anadromous and semelparous fish without homing behaviours. Despite being a freshwater, free-living organism for large part of its life cycle, its adulthood is spent as a parasite of marine vertebrates. Here, we performed the first genome-wide characterization of sea lamprey’s genetic diversity in their European natural range. The objectives were to investigate the connectivity among river basins and explore evolutionary processes mediating dispersal during the marine phase. For that, we sequenced 186 individuals from 8 locations spanning the North-eastern Atlantic coast and the North Sea with double-digest RAD-sequencing, obtaining a total of 30910 bi-allelic SNPs. Population genomic analyses reinforced the existence of a single metapopulation encompassing freshwater spawning sites within the north-eastern Atlantic and the North Sea, though the prevalence of private alleles at northern latitudes suggested some limits to the species’ dispersal. Seascape analyses revealed candidate loci associated with the abundance of some host species and were located in a genomic region coding for variable lymphocyte receptors, an adaptive immunity tool unique to jawless vertebrates, and to *MARCH* proteins, a family of E3 ubiquitin ligases also involved in the regulation of immune responses. Abiotic factors (e.g., maximum phosphate, dissolved oxygen and water temperature) were significantly correlated with candidate loci associated with the myo-inositol synthesis, a pathway linked to osmoregulation, and to other genomic regions involved in organismal homeostasis. The identification of adaptive seascapes in this ancient species, especially if linked to primitive adaptive immune responses, could be relevant to understand the evolutionary pathways early in vertebrate evolution.

## Introduction

Freshwater systems have played a key role in the evolution of human society through providing reliable sources of potable water and food, as well as communication and trading routes (Ormerod, Dobson, Hildrew, & Townsend, 2010). To facilitate these services, rivers have been heavily modified in order to facilitate navigation and provide more efficient hydropower generation through the building of, for example, weirs and dams (Dudgeon et al., 2006; Grill et al., 2019). For anadromous fishes, including some species of salmonids and lampreys, these modifications can place considerable pressure on their freshwater life phases through blocking access to environmentally stable reproductive and nursery habitats (e.g. (Branco, Segurado, Santos, Pinheiro, & Ferreira, 2012; C. S. Mateus, Rodríguez-Muñoz, Quintella, Alves, & Almeida, 2012; Moser, Almeida, King, & Pereira, 2020). Maintaining sustainable diadromous fish populations is vital both from a social-economic perspective - as many of these species are economically relevant - and from a biological standpoint, as spawning and foraging migrations ensure energy and biomass transfer between marine and freshwater environments (Limburg & Waldman, 2009).

Despite local freshwater extinctions, anadromous populations can still be maintained across large spatial scales through their use of the marine habitat, while using available rivers for spawning and nursery habitats (Moser et al., 2020). In this context, characterizing connectivity and inferring whether populations are locally adapted is essential for the sustainable replenishment of extant populations. Genetic and genomic resources have greatly contributed to understanding connectivity patterns (Asaduzzaman et al., 2020; Perrier, Le Gentil, Ravigne, Gaudin, & Salvado, 2014), defining conservation units (Catarina Sofia Mateus, Almeida, Quintella, & Alves, 2011; Waples & Lindley, 2018), and pinpointing evolutionary signatures related to migration, maturation and/or local adaptation in several anadromous species, such as in the Atlantic salmon, brown trout or shad (Antognazza et al., 2021; Kapralova et al., 2011; Leitwein, Garza, & Pearse, 2017; Prince et al., 2017). Genome-wide information is nevertheless vital to ensure the effectiveness of conservation-driven translocation activities, as it would permit to devise genetic rescue strategies to mitigate the species’ biodiversity losses and increase population resilience without compromising local adaptation (Whiteley, Fitzpatrick, Funk, & Tallmon, 2015)

The sea lamprey *Petromyzon marinus* is an anadromous, ancient jawless vertebrate whose native distribution is limited to the Western and Eastern coasts of the North Atlantic and inside the Mediterranean, where individual abundances fluctuate regionally and locally from one freshwater system to another (Guo, Andreou, & Britton, 2017; Stéphanie et al., 2020). The sea lamprey has a complex lifecycle consisting of several developmental stages, where the major morphological changes are intrinsically associated to feeding behaviour (Youson & Potter, 1979). Post-hatching larvae, or ammocoetes, are free-living organisms that feed on detritus deposited on river sediment beds or through filtration of the water column (Guo et al., 2017). The metamorphosis into young adults marks the onset of the species’ parasitic behaviour, as sea-dwelling lampreys mandatorily feed on blood of other vertebrates (Quintella et al., 2021; Silva, Servia, Vieira-Lanero, Barca, & Cobo, 2013). Adult sea lampreys spend their life at sea, a stage that is expected to last between 10 and 28 months preceding sexual maturation (Beamish, 1980; Renaud & Cochran, 2019; Silva et al., 2013). Spawning migration towards freshwater, however, is not philopatric regarding spawning locations; contrary to other anadromous fishes, sea lampreys do not return to the natal rivers to spawn (Almada et al., 2008; Bergstedt & Seelye, 1995; Spice, Goodman, Reid, & Docker, 2012). Instead, it has been suggested that sea lampreys are steered by a migratory pheromone released by stream-resident larvae (Sorensen et al., 2005; Vrieze, Bergstedt, & Sorensen, 2011), although natural recolonization of freshwater locations without ammocoete presence has been recently noted (Reid & Goodman, 2020). Less understood is how selection acts across the sea lamprey life cycle, despite its numerous life history events (such as the transition to parasitism linked to adulthood, onset of anadromous and catadromous migrations) providing substantial opportunities for selection to act. For instance, it has been recently shown that the parasitic adult stage imposes a selective pressure on the host, as suggested by evidence showing transcriptional response of genes involved on the regulation of inflammation and cellular damage in parasitized lake trout (Goetz, Smith, Goetz, & Murphy, 2016). Thus, host-parasite co-evolutionary dynamics between sea lamprey and their hosts are plausible. Indeed, recent work investigating the molecular pathway of the renin-angiotensin system on lampreys has suggested that its evolution is linked with mimicry of the teleost-like angiotensin while parasitising, in order to recreate host’s environment and avoid rejection – referred to as endocrine mimicry (Goetz et al., 2016). A putative relationship between distribution of potential hosts and signatures of selection on the sea lamprey genome - which could indicate adaptation to the host community - has not been investigated at any spatial scale.

Transitions from fresh to salt water (and vice-versa) do impose an energetic cost associated with physiological and molecular re-arrangements to cope with chemical composition of the different environments (Ferreira-Martins, Coimbra, Antunes, & Wilson, 2016). Reduced tolerance to high salinities has been reported in sea lamprey adults performing spawning migrations; failure to locate suitable spawning locations in freshwater due to connectivity loss by dams and weirs requires re-entering estuaries and the marine environment to access adjacent river systems (Sunga, Wilson, & Wilkie, 2020). Assuming that tolerance to those transitions would be an adaptive trait, it could be hypothesized that ocean salinity would play a role in limiting adult sea lamprey dispersal. Within a certain geographic area, individuals could be adapted to fresh-saltwater gradients to minimize failure of reproduction by spending time searching for suitable freshwater spawning locations. Given the regional and local extinctions of sea lamprey populations in the European range (Hansen et al., 2016; C. S. Mateus et al., 2012), the absence of homing behaviour might facilitate conservation practices based on, for example, the translocation of ammocoetes to maintain local biodiversity levels, as performed on the Columbia river basin with Pacific lampreys (Ward et al., 2012). Undertaking similar conservation measures to mitigate the local extinctions of European lamprey would thus demand an upscale of extant knowledge on the species’ connectivity and the identification of signatures associated with adaptive processes underlying their evolutionary potential (Eizaguirre & Baltazar-Soares, 2014; McMahon, Teeling, & Höglund, 2014). While traditional molecular tools have been extensively applied to investigate genetic structure or historical demography (Genner, Hillman, McHugh, Hawkins, & Lucas, 2012; C. S. Mateus et al., 2012), next-generation sequencing has been surprisingly lacking among the repertoire of studies dedicated to natural populations of sea lamprey. By harnessing the power of genome-wide information, here we intend to revisit genetic structuring among populations of sea lamprey across the North Atlantic coast while exploring molecular signatures of the selective pressures hypothesized above. Inferring signatures of selection on a highly migratory species, such as sea lamprey, would help the understanding of how microevolutionary processes may act on complex life history strategies. Concomitantly, characterizing the connectivity of spawning grounds and the evolutionary pressures that could be delimiting the marine dispersal of eastern Atlantic Sea lampreys is fundamental in establishing conservation measures befitting the species’ wide distribution range.

## Material and Methods

### Sampling strategy across hydrographic basins

To explore the evolutionary history of sea lamprey populations across the Eastern Atlantic coast, 186 individuals were sampled from eight locations across their range and located in different hydrographic regions (Fig. 1). Fin tissue samples were available from the rivers Tagus (n=20) and Mondego (n=20) in Portugal, from the Ulla (n=22) in northern Spain, from the Gironde estuary (n=18) in France, from the rivers Severn (n=22), Frome (n=34) and Yorkshire Ouse (n=20) in Great Britain and from the river Roflsan (n=22) in Sweden.

**Fig. 1.**
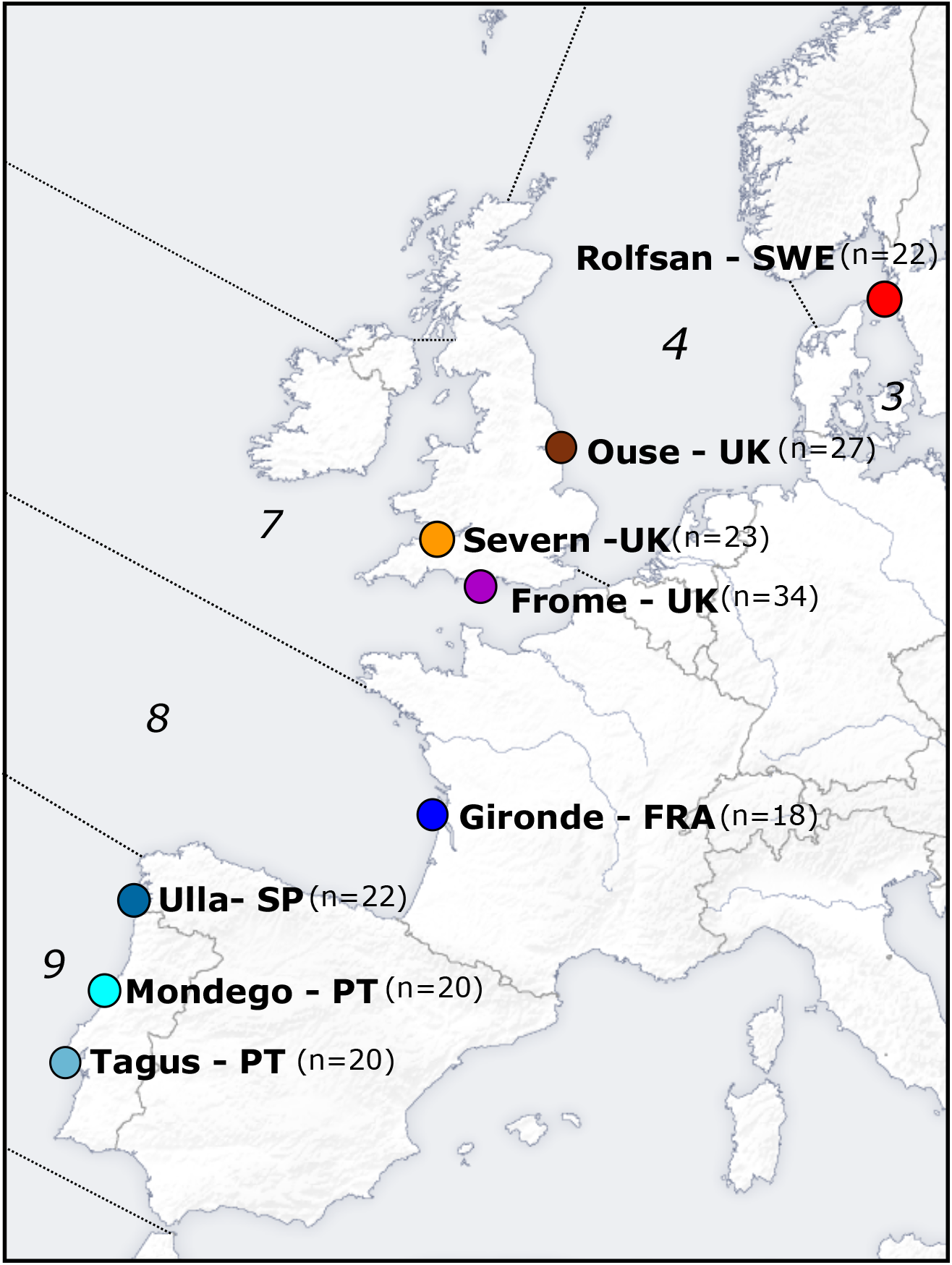
Geographic information and distribution of sampled locations. Here represented are the sampled freshwater locations (dots) and respective correspondence to FAO’s major fishing areas in the Northeast Atlantic (italicized numbers), delineated by dashed lines. Specifically, *3* corresponds to Skagerrak, Kattegat, Sound, Belt Sea, and Baltic Sea, *4* to the North Sea, *7* to Irish Sea, West of Ireland, Porcupine Bank, Eastern and Western English Channel, *8* to the Bay of Biscay and *9* to the Portuguese waters. In parenthesis, the number of individuals collected for each location utilized in this study.

Migrating sea lamprey were captured using unbaited two-funnel eel pots. Captured individuals were held in water-filled containers (100 L) prior to general anaesthesia (MS-222) that enabled a fin biopsy to be taken. All the sampled individuals were adults, except for those from the Frome, where, due to unavailability of adults during sampling period, ammocoetes were sampled. Ammocoete sampling was performed by dredging, with a dipnet, soft sediment areas of the river Frome. All of the tissue samples were preserved in ethanol (96 %) prior to extraction.

### DNA extraction, library preparation and double-digest Restriction-Associated DNA tag sequencing

Genomic DNA was extracted from the fin or muscle tissues with Qiagen DNeasy Blood and Tissue kit (Hilden, Germany) following the manufacturer’s instructions. Library preparation, sequencing and bio-informatic processing of raw reads (demultiplexing of individual barcodes, removal of adaptors and barcodes and trimming) was performed at the GENEWIZ© in Takeley, United Kingdom. Library preparation followed the double digest protocol of Peterson et al. (2012). Libraries were constructed by pooling 48 samples after individual barcoding. Genomic DNA was digested with EcoR1/Msp1 restriction enzymes. Two 96-well plates were sequenced on an Illumina HiSeq2500 with a 2×125bp paired-end (PE) configuration. Final library was size-selected between 300-400bp inserts. Only reads with Q>30 was kept for downstream analysis.

### Processing paired end reads

Reads were aligned against the available sea lamprey genome (Smith et al., 2013) using the *bwa-mem* algorithm (Heng Li, 2013) implemented in *bwa* (H. Li & Durbin, 2009). Bam files were imported into Stacks, version 2.2 (Catchen, Hohenlohe, Bassham, Amores, & Cresko, 2013) for further analyses and variant calling. Only bi-allelic loci that were present in 80 % of the populations were kept, as well a one SNP per tag (Rochette & Catchen, 2017). Because the sampling spanned a wide geographic area, minor allelic frequency of 0.01 was filtered out to avoid losing unique information. To mitigate a potential effect of the high repeatability found on this species’ genome, all the reads whose coverage was above 100x were also filtered out, along with loci that mapped to more than one catalogue.

### Genetic diversity indices across sampled locations and hydrographic regions

All statistical analyses were computed in R v 3.4.1. (R Core, 2013). For each location, we estimated the heterozygosity (*Hs*), and inbreeding coefficient (F) with the *adegenet* R package (Jombart, 2008) and the number of private alleles (*pA*), nucleotide diversity (pi) with *Stacks*. Absolute values of *pA* were log transformed to normalize the distribution prior to statistical analyses. The distribution of genome-wide diversity was first compared across sampled locations, where we investigated the relationship of each index with latitude and longitude. Relationships were tested between diversity indices and geographic distance using linear models.

### Population structure and connectivity among freshwater locations

Multiple approaches were used to investigate the population connectivity between the sampled locations. A PCA was performed in the R package *pcadapt* v 4 (Privé, Luu, Vilhjálmsson, & Blum, 2020) in a first attempt to verify if sampling locations harboured genetically similar individuals. Pairwise F_ST_ was estimated in Arlequin v3.5 (10000 permutations) (Excoffier & Lischer, 2010) to understand overall genome-wide diversity. Bayesian clustering methodology explored the distribution of genome-wide variation across the sampled sites, running *faststructure* (Raj, Stephens, & Pritchard, 2014) with three independent iterations for *K* values of 1 to 8. The most likely *K* number was assessed using *chooseK* (T. Raj et al., 2014). Admixture proportions were visualized with *compoplot* function in the *adegenet*. Lastly, discriminant analysis of principal component (DAPC) was used to assess individual clustering based on genetic similarities in *adegenet*. We performed DAPCs assuming both the original sampling location, i.e. predefined populations as well as exploring configuration of clusters with the function *find.clusters*. Putative clusters were identified with Bayesian Information Criteria (BIC) and assignment of individuals was compared to that of sampling locations.

### Investigating the relationship between genomic variation and environmental variables

Environmental and genetic variation tend to be interpreted together under the assumption that co-variation suggest signatures of evolutionary responses to environmental conditions (Günther & Coop, 2013). The wide range of environments experienced by sea lampreys during their life cycle renders environmental correlations a logical approach to investigate putative signatures of adaptation across spatial scales. Specifically, we investigated possible adaptations to the transition freshwater-saltwater-freshwater by investigating the relationship between genomic variation *versus* sea surface salinity. Adaptation to the nearby marine environment was further investigated by inspecting the relationships between genomic variation with surface temperature, a surrogate for water temperature, and dissolved oxygen concentration, both critical for ectotherm thermal regulation and metabolic processes in the marine environment. We also explored whether coastal anthropogenic activities could be a mediating factor to sea lamprey’s marine distribution. We considered phosphate and nitrate concentrations as they are commonly associated with sewage effluent discharges and the consequent eutrophication of coastal systems at various spatial resolutions (Van Drecht, Bouwman, Harrison, & Knoop, 2009).

All these variables were extracted from Bio-ORACLE (Assis et al., 2018; Tyberghein et al., 2012) with customized R scripts. A total of 6 environmental parameters, i.e., *sea surface salinity, sea surface temperature, dissolved oxygen, phosphate concentration* and *nitrate concentration* were collated, including the minimum and maximum of the variables described in Table S1.

The obligatory parasitic stage of sea lamprey’s life cycle could suggest the species might have evolved adaptations towards availability of certain host species or to the diversity of host communities. We tested that hypothesis by comparing the variation in the abundance of fishes that have been referenced in the literature as bearing sea lamprey sucking marks or adult individuals attached to them, to the genome-wide loci variance among sampled locations. The list of potential hosts included the Atlantic mackerel (*Scomber scombrus*), the Atlantic cod (*Gadus morhua*), the basking shark (*Cetorhinus maximus*), the European hake (*Merluccius merluccius*), the thick lip grey mullet (*Chelon labrosus*), the Atlantic salmon (*Salmo salar*), the twaite shad (*Alosa fallax*) and the allis shad (*Alosa alosa*) (Silva, Araújo, Bao, Mucientes, & Cobo, 2014). Abundances were extracted from the Food and Agriculture Organization (FAO)’s FishStatJ - Software (FAO, 2016) as capture data (in tonnes) (Fig. S1). We focused on areas within the Northeast Atlantic (Major Fishing Area 27) which corresponded to marine regions adjacent to sampled sites. More specifically, we collected data from area 3, corresponding to Skagerrak, Kattegat, Sound, Belt Sea, and Baltic Sea; area 4, the North Sea; area 7, corresponding to Irish Sea, West of Ireland, Porcupine Bank, Eastern and Western English Channel; area 8, the Bay of Biscay; and area 9, Portuguese waters. Captures on FishStatJ spanned a period of 10 years from 2006 to 2016, thus we calculated the 10-year harmonic average for each species, per area. To reduce the variance associated with annual or decadal means, all abiotic and biotic variables were standardized for their z-scores prior to computation analyses. Standardized values were further used throughout the remaining analyses.

The relationships between the above mentioned variables and genetic variation was first tested for the full SNP dataset in Bayenv2 (Günther & Coop, 2013). To increase the robustness of any observed gene-environment association, we imposed a double filter for statistical significance. Specifically, only the gene-environment relationships with a Bayes Factor > 10 (Jeffreys, 1998) and an Spearman’s *rho* statistic above 0.2 (Günther & Coop, 2013) were considered as significant. In addition, we utilized latent factor mixed models (LFMM) to further inspect for environment correlations with the genetic data (Frichot, Schoville, Bouchard, & François, 2013). Contrary to the island-model of neutral allelic frequency distribution to identify outliers assumed in Bayenv2, LFMMs require *a priori* information on population structure to infer the background noise of neutral differentiation (Frichot et al., 2013). We ran LFMM on the R package *LEA* (Frichot & François, 2015). Because the methodology is sensitive to missing data, we performed imputation of null alleles based on estimated ancestry coefficients and ancestral genotype frequencies, as suggested by Frichot & François (2015) and considering the “mode” of each allele. We set the K to assume all possible populations, i.e., 1 to 8, we replicated each run 10 times, and applied a random set of 3000 SNPs to initialize the cross-entropy algorithm. Outliers were identified considering a z-score distribution of correlation values after p-value adjustment, as suggested by Frichot et al (2013). This approach has been shown to be potentially complementary to other correlation-based strategies, though with a higher than average number of positives (Nielsen, Henriques, Beger, Toonen, & von der Heyden, 2020). As such, we further controlled for false discovery rate (FDR) according to Benjamini-Hochberg procedure with a q = 0.1 (Benjamini & Hochberg, 1995). Lastly, we utilized *pcadapt* to screen for outlier loci. The number of PCs to retain was chosen accordingly to the scree plot curve and p-values were subject to Bonferroni correction with α = 0.001.

Only loci identified as outliers in all detection methods were classified as candidates under selection and were used in further analyses. We calculated diversity estimates and inferred population structure with the same methodology applied to the full loci dataset.

### Candidate loci annotation

Genomic sequences flanking candidate loci were retrieved from the sea lamprey genome (P.marinus_7.0) at Ensembl’s database by extending the size of the RADtag 1000bp upstream and downstream the coordinates to which they mapped against. Extending the RADtag alongside the reference genome was performed with the intent of capturing regions potentially linked to the candidate loci. Fragments were blasted against the NCBI database.

## Results

### Sequencing statistics and filtering outputs

Overall, mapping resulted in a mean percentage of 36 % (SE = 0.3%) reads kept, representing a mean of 1.94 × 10^6^ (SE=596.66) reads per individual. Mean coverage per individual was 20.25x (SE = 0.79), with the mean number of genotyped sites per locus being 269.83 (SE = 0.24); 30910 loci were retained for downstream analysis.

### Genetic diversity estimates

Genetic diversity estimates were similar among sampled locations when considering all sequenced sites, with *Hs* varying between 0.10 for the Frome 0.12 for other 6 locations (Table 1). The number of private alleles had higher variation, with the highest number found in the Rolfsan (228) and the lowest in Gironde (77), with the difference being significant according to geographic distance (R^2^=0.18, p=0.01) (Fig. 2a), where the higher the distance between sampling areas, the greater the differential of private allele numbers between the sampling areas. There was also a significant and positive linear relationship between the number of private alleles and latitude (R^2^=0.72, p=0.04) (Fig. 2b), suggesting an overall pattern of increased unique genomic diversity at higher latitudes.

**Table 1.**
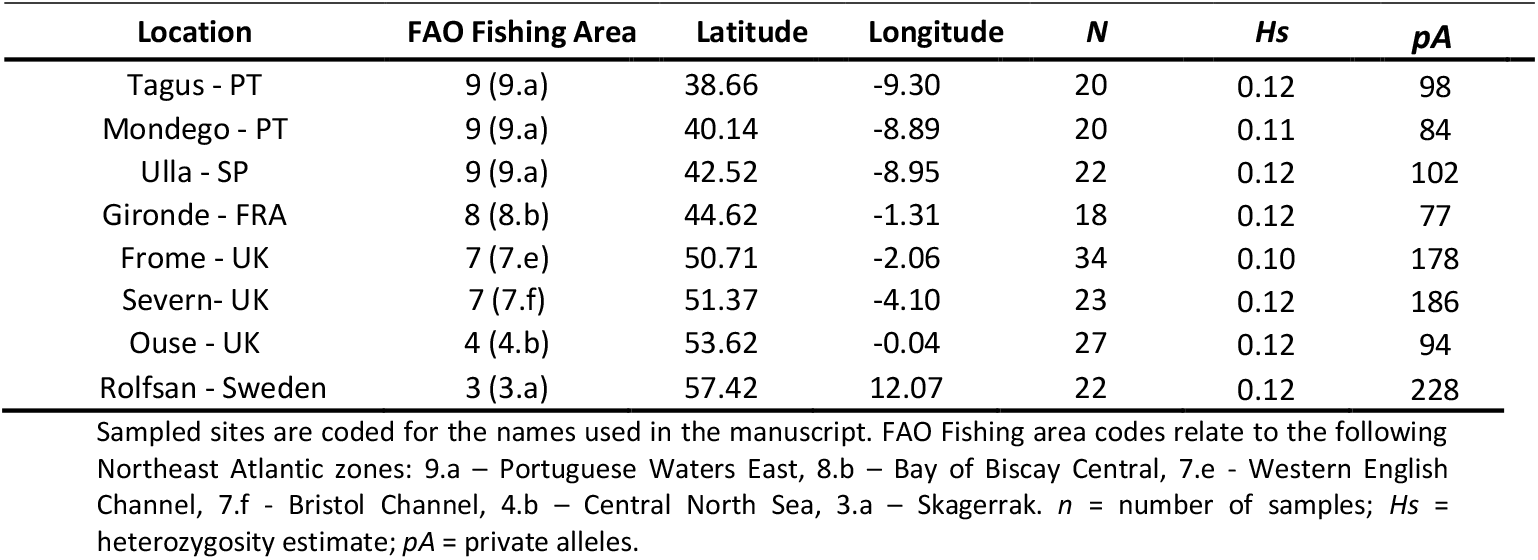
Geographic information and diversity indices of sampled locations.

**Fig. 2.**
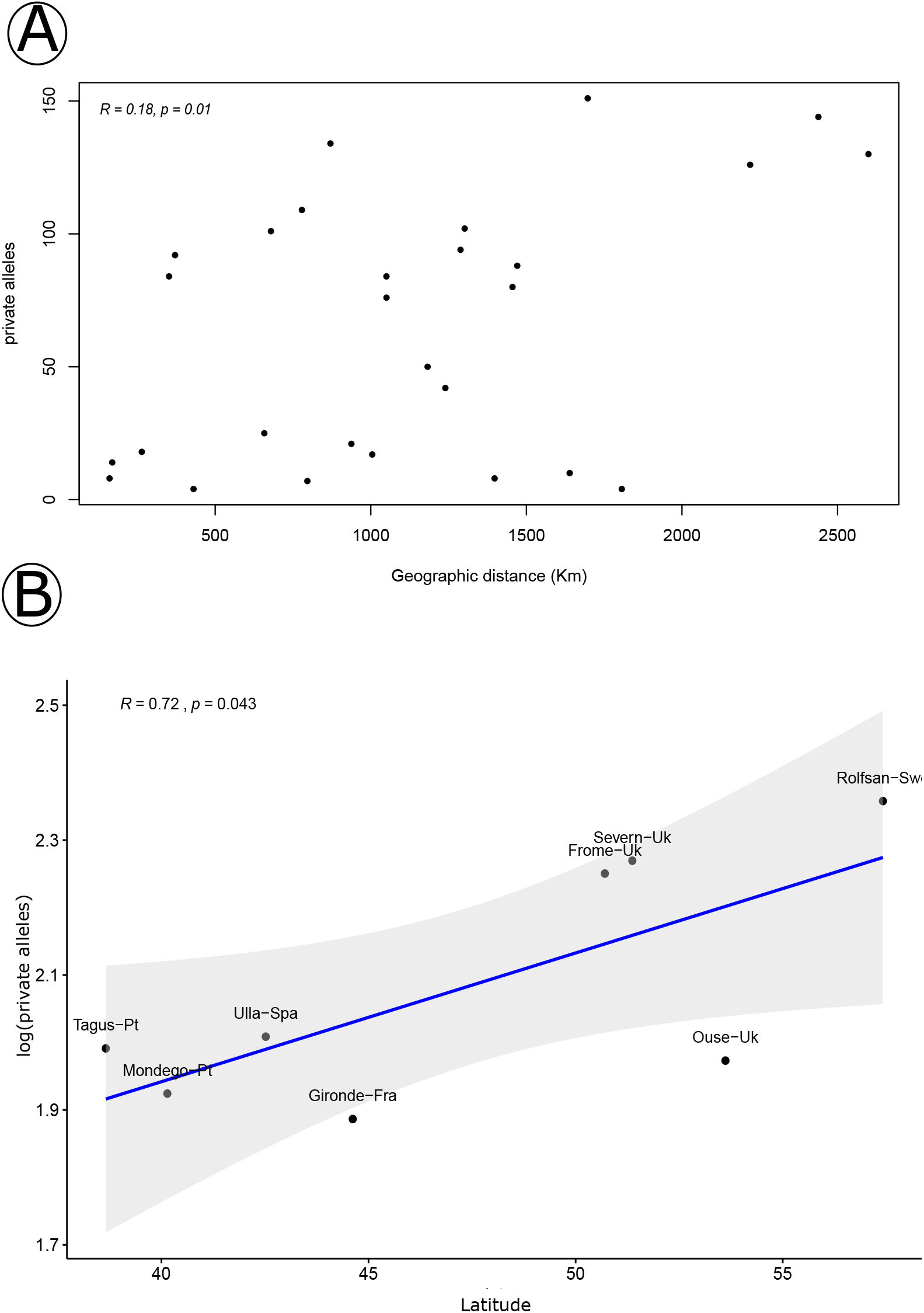
Relationships between location-specific alleles and geography. Panel A represents the relationship between the absolute number of private alleles differences (x-axis) as a function of geographic distance (y-axis) between population pairs. Panel B investigates the relationship between latitude (x-axis) and the log-scaled number of private alleles (y-axis) with sampled locations marked in the dots. Smooth bands of the linear model log(pa)~latitude was applied to aid the visualization of the trends.

### Population structure and connectivity among freshwater locations

A principal component analyses revealed some individuals associated with the locations of Frome and Rolfsan to be separated from the cloud of points representing all other samples (Fig S1). For the DAPC, 150 PCs were retained that explained 85% of variation of the overall SNP dataset. Exploring the number of clusters (1-8) with *find.cluster* function and subsequent ordinate plotting and considering function-defined groups, we observed that assignment did not match sampling locations(Fig. S2); when we defined grouping concordant with sampling locations, we observed a clearly distinct separation between Rolfsan and all other individuals (Fig. 3). Pairwise F_ST_ values were also generally low and non-significant, except between the Rivers Frome and Rolfsan (F_ST_ = 0.010, p = 0.001), and Frome and Ouse (F_ST_ = 0.006, p = 0.001) Table S2).

**Fig. 3.**
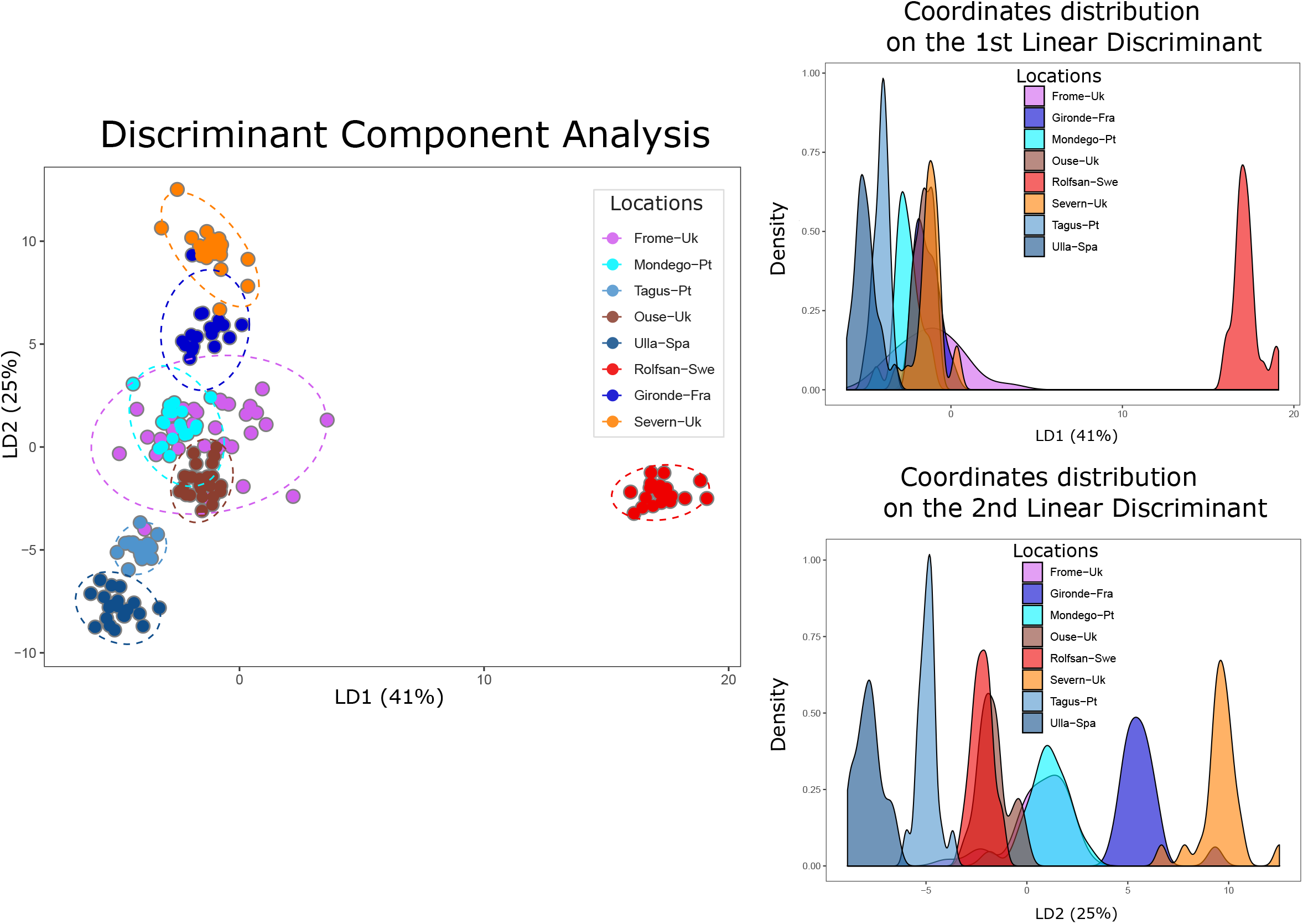
Discriminant Analyses of Principal Components (DAPC) with pre-defined population groups. Abstract distribution of individuals according to linear discriminant functions (1 and 2) that together captured 66% of genome-wide variation. Ellipses in dashed lines represent the 95% confidence interval of a normal distribution of PCs.

### Gene-environment associations and gene ontology of putative candidate loci

There were 118 loci with a Bayes factor > 10; amongst these, 42 also had an absolute value of *rho* > 0.20. The *pcadapt* approach, which accounts for population differentiation while identifying outliers, detected 629 candidate loci across variables. LFMMs detected candidate loci, per variable, in the range of those detected by *pcadapt*. However, decreasing the adopted *q* (from the suggested *q*=0.1 to *q*=10^−3^) to adjust for false discovery rate did not significantly reduce putative outliers. Instead, it placed their number in the range of what was observed by Nielsen et al (2020), i.e., 1/5^th^ of SNPs in the dataset were deemed outliers. Altogether a total of 11 unique candidate loci was shared among approaches, which the LFMM analyses distributed as following: *maximum sea surface temperature, minimum nitrogen concentration, maximum dissolved oxygen, cod abundance* and *twaite shad abundance* correlated with 1 locus each; *maximum phosphorus concentration, minimum phosphorus concentration* and *basking shark abundance* correlated with 2 loci while 3 loci *allis shad abundance* (Fig. 4). Heterozygosity estimates of candidate loci within populations were substantially above those considering the full panel of SNPs and ranged between 0.11 in Ulla and 0.20 in Mondego (t-test: *Hs*_mean_ All = 0.12; *Hs*_mean_ candidate= 0.16; p = 0.002). We did not detect population structure associated with candidate loci allelic frequencies: overall F_ST_ of −0.001 and the non-identification of clusters either with or without *a prior* population information inserted into DAPCs (Fig. S4).

**Fig. 4.**
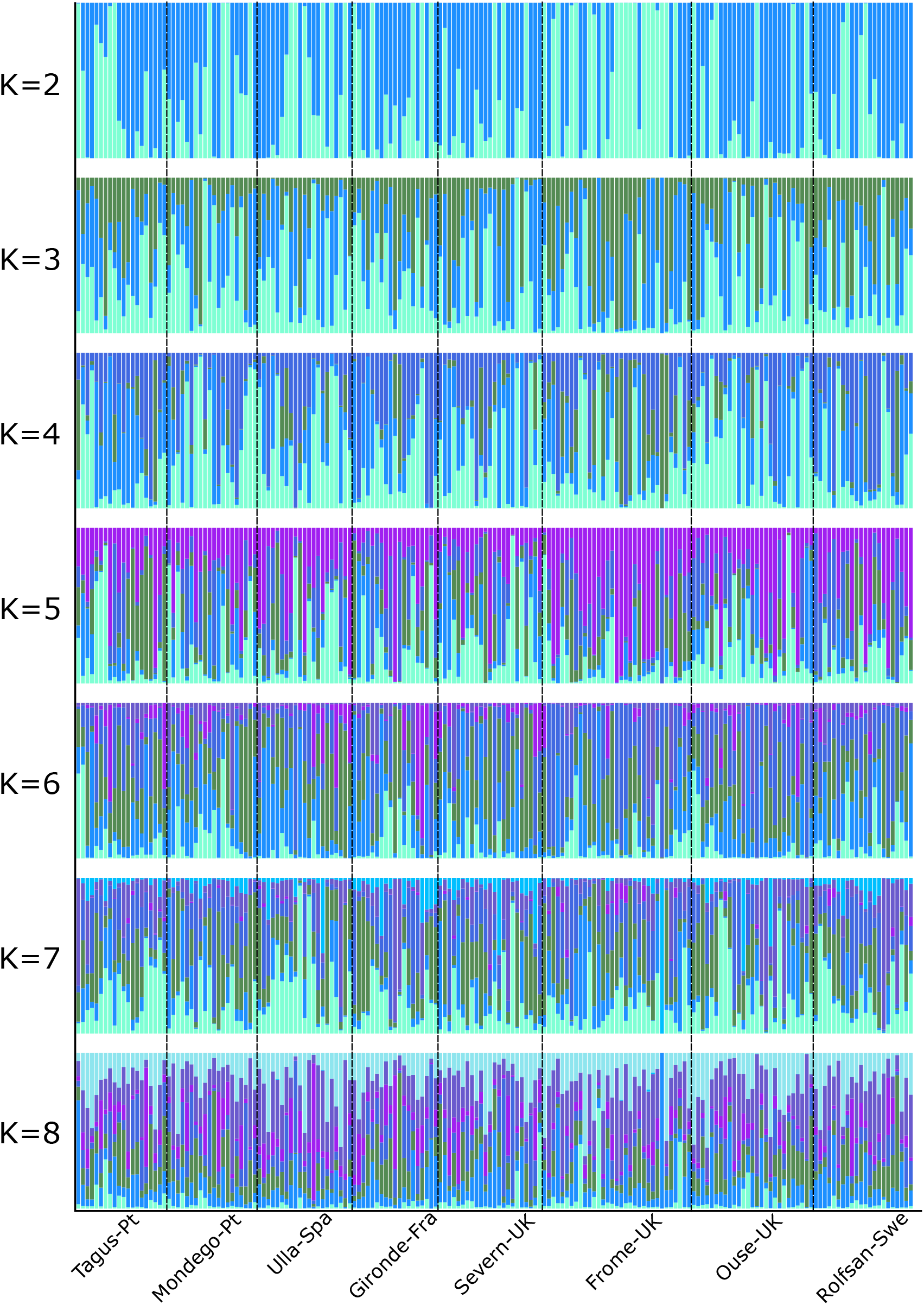
Bayesian clustering to explore number of K in relation to sampling locations. Bayesian clustering generally suggests an high homogeny of admixture probabilities (y-axis) among sampling locations (x-axis), a pattern reproducible independently of the explored number of Ks.

**Fig. 5.**
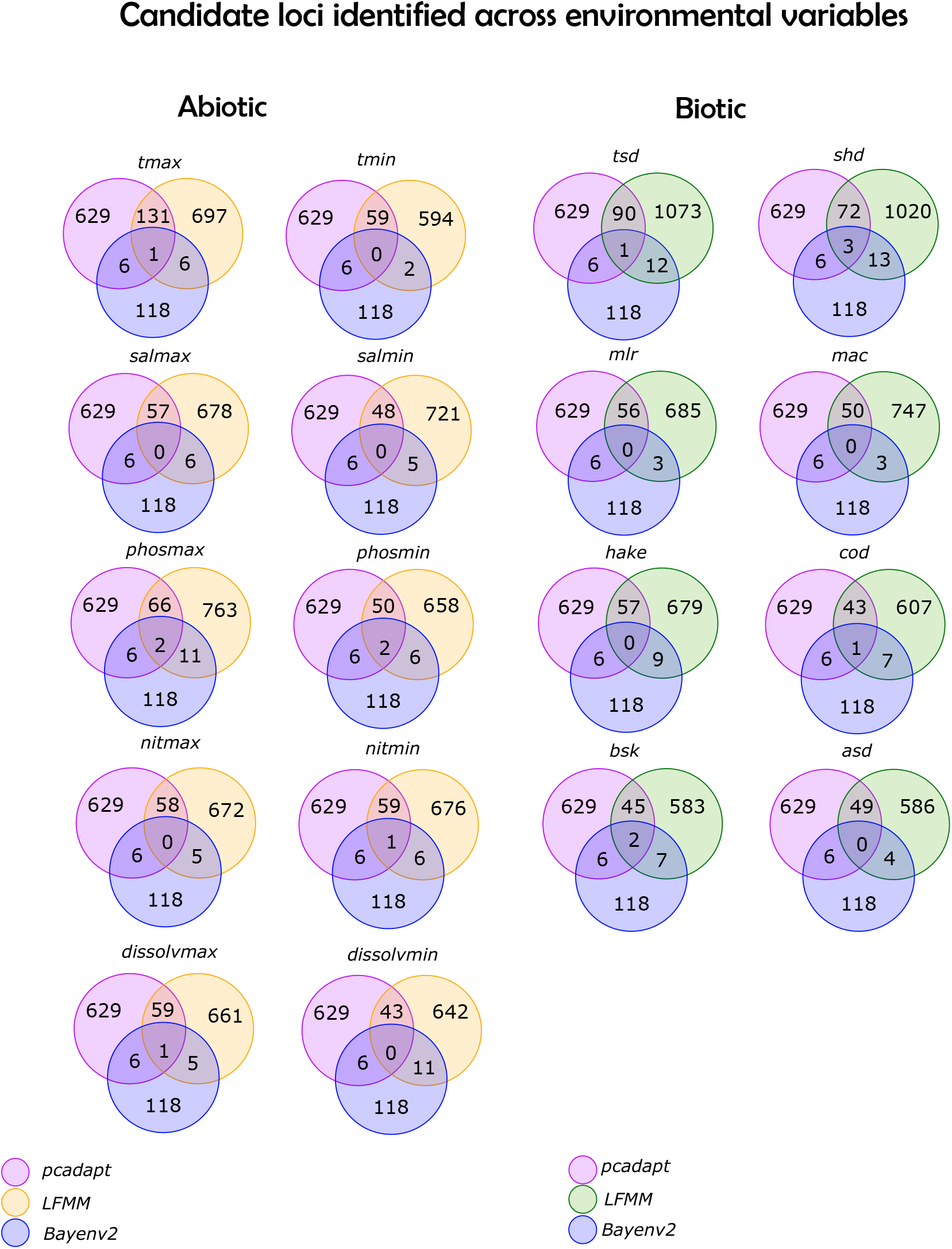
Venn diagram of candidate loci identified by multiple outlier detection techniques. Shown here are the number of candidate loci identified by each software. LFMM approach implemented in the R package *lea* is coloured in yellow for the abiotic variables and in green for the biotic variables.

Blasting the 11 candidate loci against NCBI database revealed interesting patterns in light of the selective pressure against which their frequencies relate. For instance, candidate loci associated with biotic variables *allis shad abundance* and *Atlantic cod abundance* (1838 and 13235) were found to be located in the genomic regions corresponding to the *jVLR* gene encoding for variable lymphocyte receptors, which allow lymphocytes to detect antigens in their environment (J. Li, Das, Herrin, Hirano, & Cooper, 2013). Locus 1838 also aligned against the SAFB-like transcription modulator, the *NF-kB* gene and the Hox3 homeobox protein, features involved in transcription regulation and RNA binding, and cellular response to stress. Candidate 13235 was also identified in a region of the sea lamprey genome’s Zinc finger OZF-like protein. Lastly, locus 13283 identified as a candidate to *twaite shad abundance* to a membrane associated RING-CH type 3 finger ubiquitin ligase and critical regulators of immune responses (Lin, Li, & Shu, 2019).

In relation to the putative abiotic pressures, candidate locus 26013 was associated with both *maximum phosphate concentration* and *maximum dissolved oxygen concentration* and mapped to a genomic region coding identified as the par-3 family cell polarity regulator beta (*pardb3* gene), a highly conserved feature across metazoan genomes involved in cell polarity and adhesion (Thompson, 2022); loci 13513 and 14785, which associated with *maximum phosphate concentration* and *maximum temperature* respectively, mapped against *P. marinus’* inositol-3-phosphate synthase 1 (*isyna-1* gene) a molecule involved in the production of myo-inositol in a wide range of organisms including fish (Agranoff, 2009) and to *P. marinus’* homeodomain *HOXW10A-like* gene and Tcl-1 like transposon, both previously identified to be associated with long-term plasticity of spinal cord locomotor network in sea lamprey (Bevan, Vakharia, & Parker, 2008). Loci 18413 and 27929 were mapped against highly repetitive microsatellite regions (Table 2).

**Table 2.**
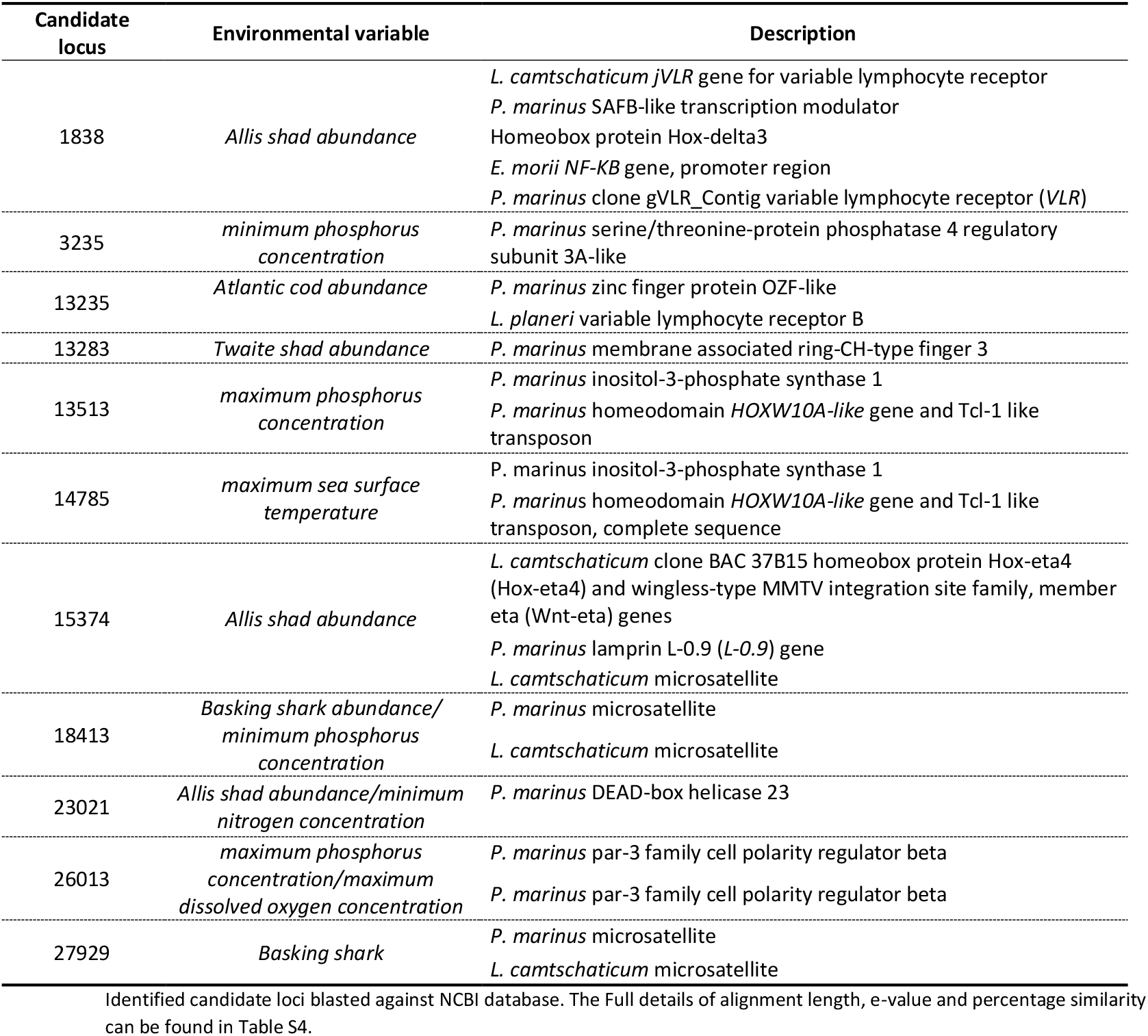
Annotations of candidate loci against NCBI database.

## Discussion

Despite being a key species in the understanding of the evolution of vertebrates (Smith et al., 2013; Sower, 2018), there remains a paucity of information on how populations of sea lamprey evolved in natural habitats. With their abundance declining in freshwater locations alongside the eastern Atlantic shore (Hansen et al., 2016), understanding connectivity among spawning locations, as well as the evolutionary potential of the species, is a pressing issue towards conservation of this ancient organism. Here, we present the first genome-wide screen of sea lampreys’ genetic diversity, highlighting the microevolutionary processes governing the species’ limited dispersal and to shed light on early stages of vertebrate evolution.

### Sea lampreys constitute a large Eastern Atlantic metapopulation limited by northward dispersal

The absence of homing behaviour towards natal spawning grounds renders sea lamprey an exception among diadromous species. Thus, the migratory behaviour associated with the mandatory marine phase of sea lamprey would suggest a homogenisation of genetic variance across the species’ distribution range. It is not surprising then that patterns of panmixia have been reported among populations of sea lamprey on the western Atlantic coast (Waldman, Grunwald, & Wirgin, 2008), on the eastern Atlantic coast (Almada et al., 2008; Lança et al., 2014; C. S. Mateus et al., 2012; Rodríguez-Muñoz, Waldman, Grunwald, Roy, & Wirgin, 2004), as well as among populations of Pacific lamprey *Entosphenus tridentatus* (Spice et al., 2012). Despite the upscale to a genome-wide representation of molecular signatures, we observed a strikingly similar pattern of panmixia supported both by homogenised genetic diversity and almost complete absence of differentiation among sampled locations. However, our results did suggest that Atlantic sea lampreys may have limits to their dispersal, as also identified in their Pacific counterpart (Spice et al., 2012). Support of a limited dispersal scenario was provided by both the discriminant analyses on the whole SNP dataset, which pinpointed the north-easternmost location of Rolfsan with some degree of differentiation in relation to all other locations, as well as a significant and positive correlation between the number of private alleles and latitude, which may be interpreted as a signature of limited connectivity with southern populations. The barrier imposed by the salinity gradient to other marine species across the Kattegat-Skagerrak region which defines the transition zone between the North-East Atlantic and the Baltic Sea (Johannesson, Le Moan, Perini, & André, 2020; Wennerström et al., 2013), is perhaps the most parsimonious explanation for the subtle differentiation and the high number of private alleles in the Rolfsan. In this context, the Rolfsan’s lampreys might be part of a sub-population more likely to spawn within the Baltic Sea area, a scenario supported by scarce modern-era observations of sea lamprey spawning aggregations in the Vistula river, Bay of Gdansk and Szczecin lagoon, but not in German rivers (Thiel et al., 2009).

### Investigating dispersal limitations under the magnifying-glass of molecular data

With generally low genome-wide differentiation over large spatial areas, allelic frequencies deviating from the neutral spectrum in marine populations have been increasingly associated with environmental cues characteristic of a heterogenous oceanic environment (P.-A. Gagnaire & Gaggiotti, 2016). For example, seascape genomics have revealed that despite high levels of gene flow, temperature could explain distribution patterns in abalone (*Haliotis laevigata*), lobsters (Benestan et al., 2016) and cod (Clucas, Lou, Therkildsen, & Kovach, 2019; Therkildsen et al., 2013), while salinity extended associations to herring and cod (Berg et al., 2015; B. Guo, Li, & Merilä, 2016). Sea lampreys have a very specialised set of life-history traits whereby the freshwater phase hinders a full-scale open ocean dispersal, as shown by the stark differentiation between Eastern and Western Atlantic coasts (Genner et al., 2012), and their obligate parasitism could render their dispersal to be passively dependent on that of the hosts. Thus, it has been argued that limits to marine dispersal might be imposed both by abiotic and biotic factors (Guo et al., 2017). The routine use of outlier detection methods and/or environmental associations (genotype-environment associations – GEA) have assisted the exposure and identification of suitable analytical strategies to overcome issues related to high frequencies of false positives (type II errors), known to be inherent to analytical processes, algorithms, and sampling strategies (Capblancq, Luu, Blum, & Bazin, 2018; Lotterhos & Whitlock, 2015; Narum & Hess, 2011; Nielsen et al., 2020).

By utilizing three different approaches to identify candidate loci under selection, we expected to mitigate issues related to high frequencies of false positives (type II errors), known to be inherent to analytical processes, algorithms, and sampling strategies (Capblancq et al., 2018; Lotterhos & Whitlock, 2015; Narum & Hess, 2011; Nielsen et al., 2020). Here, we report a relationship between the abundance of confirmed host species’ such as the Atlantic cod, and the twaite shad, and allelic frequencies of loci located within the vicinity (maximum 1000bp) of genomic regions annotated to immune-related functions. Notably, variable lymphocyte receptors are an immune tool specific to jawless vertebrates, and RING-CH-type finger (MARCH) protein, is part of a family of E3 ubiquitin ligases also involved in the regulation of immune responses (Lin et al., 2019). Oher genomic regions identified either on these loci or in other loci associated with these same biotic variables further support regulatory functions to be under selection in our target species. These results are suggestive of a hypothetical scenario of local adaptation where sea lampreys have evolved immune responses against the antigens of the most likely hosts that they will encounter at sea. It is known that sea lampreys elicit immune response from their hosts (Bullingham, Firkus, Goetz, Murphy, & Alderman, 2022; Goetz et al., 2016), and thus it is plausible, at the light of host-parasite coevolution, to expect adaptive signatures on the sea lamprey genome (Ebert & Fields, 2020; Eizaguirre, Lenz, Kalbe, & Milinski, 2012). To the best of our knowledge, this is the first time that genomic signatures of putative adaptation to host species has been shown in sea lamprey and consequently, at such an early stage of vertebrate adaptive immune system evolution (Boehm et al., 2012; Finstad & Good, 1964; J. Li et al., 2013). A plausible caveat to these observations relies on the accuracy of host abundances collected for this study. Food and Agricultural Organization fisheries datasets are based on capture and aquaculture data, and despite being extensively utilized to understand trends in fish abundance in the context of sustainable fisheries (Froese, Zeller, Kleisner, & Pauly, 2012; Garibaldi, 2012; Naylor et al., 2021), their interpretation has historically experienced criticism (Daan, Gislason, Pope, & Rice, 2011; Pauly & Zeller, 2016). Alternatively, it could be the case that putative adaptive signatures reflect host-parasite co-evolution where the sea lamprey is the host. However, such scenario would imply the frequency distribution of a hypothetical sea lamprey parasite to co-vary with abundance of sea lamprey hosts (here investigated) across spatial scales. As in the case of sea lamprey being the parasite, data on sea lamprey’s parasites is scarce and the functional link to fitness remains elusive (Shavalier, Faisal, Moser, & Loch, 2021).

From an abiotic perspective, we find it interesting that loci associated with *maximum phosphate concentration* and *maximum temperature* relate to a genomic region coding for the *isyna-1* gene, involved in the myo-inositol (formerly referred to as vitamin B8) pathway. The biosynthesis of myo-inositol is an evolutionary conserved pathway (Majumder, Chatterjee, Dastidar, & Majee, 2003), known to modulate several vital physiological functions in living organisms, such as growth, immune efficiency, osmoregulation and exposure to heavy metal concentrations (Cui et al., 2022). Interestingly, this locus has been identified as candidate under selection also in the sand goby (*Pomatoschistus minutus*) in the context of adaptations to less saline conditions after post-glacial expansion (Leder et al., 2021). Because the same candidate loci also identified other genomic features such as the *HOXW10A-like* gene and the Tcl-1 like transposon (both involved in highly conserved mechanisms of homeostasis and plasticity of spinal cord locomotor network, respectively) it is suggested that the selection for organismal balance is plausible in the context of the heterogenous environment that sea lamprey experiences during its life cycle. While the fresh- to saltwater transition is generally important, our results suggest also the importance of the abiotic environment and putative anthropogenic impacts, such as phosphate concentration (Farmer, 2018), which along with dissolved oxygen concentration, is critical for the thermal regulation and metabolic processes of aquatic ectotherms (H. O. Pörtner & Knust, 2007). Temperature has also been shown on multiple times to be a selective pressure in the marine environment (Conover, Clarke, Munch, & Wagner, 2006; H.-O. Pörtner et al., 2001). Nevertheless, we are unable to provide evidence to support the functionality of the associations reported in our study. In a broader context, we argue that these patterns are likely to relate to spatially varying selection, a scenario previously reported in the American eel (*Anguilla rostrata*), a catadromous species with an equally diverse distribution range and complex life cycle (P. A. Gagnaire, Normandeau, Cote, Hansen, & Bernatchez, 2012).

### Concluding remarks

We show that sea lampreys in the Eastern Atlantic coast constitute a nearly panmictic population, with North Sea imposing some limits to the species’ dispersal capacity. Within the limitations of our sampling strategy, we support the idea that the North Sea-Baltic Sea transition can be a reservoir of unique sea lamprey diversity (whether it be genetic, life history-based, etc), and thus monitoring and conservation of this anadromous species should be integrated into activities that aim to preserve biodiversity in this region. Regarding the factors mediating the dispersal ability of a species without homing behaviour, we showed one possibility would be spatially varying selection shaping the adaptive seascape across sea lamprey distribution. While we find particularly interesting the association between candidate loci mapping to jawless-specific immune genes and host abundances, we acknowledge more work is necessary to further validate these results. Nevertheless, these observations could provide the background to investigate how host-parasite co-evolution could have mediated the evolution of earlier versions of vertebrate adaptive immune repertoire under natural contemporary conditions.

## Acknowledgments

This work was supported by a MSCA-IF (ADAPTATION) attributed to MBS. CSM is supported by National Funds through FCT (Foundation for Science and Technology) through the project “EVOLAMP - Genomic footprints of the evolution of alternative life histories in lampreys” (PTDC/BIA-EVL/30695/2017). CSM, PRA, BRQ and MBS are supported by the FCT strategic plan for MARE (Marine and Environmental Sciences Centre) under Project UIDB/04292/2020. Maria C. Rodicio, Mario Lepage and Micael Söderman for donation of samples.

## Data archiving statement

To be completed after manuscript is accepted for publication.

## Supplementary material

**Fig. S1 – Host community, based on capture data, across FAO fishing zones**

Cumulative histograms of averaged host captures across the FAO zones on which sampled sites were located

**Fig. S2 – Principal component analyses**

Exploring population structure with PCA.

**Fig. S3 – Discriminant analyses of principal components with full dataset**

Exploring how the number of putative clusters influences sample distribution in the abstract space and possible relationships to sampling locations utilizing full set of SNPs.

**Fig. S4 – Discriminant analyses of principal components for candidate loci**

Exploring how the number of putative clusters influences sample distribution in the abstract space and possible relationships to sampling locations utilizing only candidate loci that overlap across outlier identification techniques.

## Notes

### Competing Interest Statement

The authors have declared no competing interest.

