## Supplementary Tables for "Seascape genomics reveals limited dispersal and suggest spatially varying selection among European populations of sea lamprey (*Petromyzon marinus*)"

| **Population** | **max DO** | **min DO** | **max Pho** | **min Pho** | **max SST** | **min SST** | **max SSS** | **min SSS** |
| --- | --- | --- | --- | --- | --- | --- | --- | --- |
| Frome | 307.881 | 240.930 | 0.348 | 0.160 | 18.060 | 6.531 | 35.187 | 33.350 |
| Gironde | 292.486 | 222.956 | 0.069 | 0.000 | 21.814 | 10.286 | 34.869 | 28.641 |
| Mondego | 270.577 | 229.094 | 0.195 | 0.007 | 20.250 | 12.850 | 35.746 | 33.503 |
| Ouse | 324.894 | 239.261 | 0.630 | 0.213 | 15.543 | 4.247 | 34.504 | 33.603 |
| Rolfsan | 392.700 | 252.160 | 0.258 | 0.069 | 19.974 | -0.298 | 30.111 | 14.269 |
| Severn | 319.043 | 247.024 | 0.377 | 0.002 | 17.780 | 6.490 | 33.617 | 30.819 |
| Tagus | 265.619 | 227.363 | 0.130 | 0.033 | 20.300 | 13.746 | 35.930 | 34.150 |
| Ulla | 275.732 | 231.313 | 0.255 | 0.002 | 18.785 | 12.148 | 35.800 | 33.500 |

Table S1 – Raw abiotic values extracted from Bio-Oracle database., DO = Dissolved oxygen, Pho= Phosphate concentration SST = sea surface temperature, SSS= sea surface salinity

| **Population** | **max NIT** | **min NIT** | **DD** |
| --- | --- | --- | --- |
| Frome | 1.149 | 0.037 | 1.700 |
| Gironde | 8.245 | 0.436 | 650.000 |
| Mondego | 1.366 | 0.005 | 108.300 |
| Ouse | 3.574 | 0.023 | 51.700 |
| Rolfsan | 1.569 | 0.072 | 5.000 |
| Severn | 4.215 | 0.208 | 61.170 |
| Tagus | 0.653 | 0.001 | 500.000 |
| Ulla | 2.936 | 0.019 | 79.300 |

Table S1 (cont.) NIT = Nitrate concentration, DD= River discharge rate

Table S1(cont.) – Landings of potential sea lamprey hosts on the FAO-defined area for each of the location. Units are in tonnes

| **Population** | **Mackerel** | **Grey mullet** | **Salmon** | **T. shad** | **A. shad** | **Basking shark** | **Cod** | **Hake** |
| --- | --- | --- | --- | --- | --- | --- | --- | --- |
| Frome | 1003.194 | 14.447 | 0.308 | 0.078 | 0.671 | 0.004 | 126.309 | 608.963 |
| Gironde | 1085.738 | 39.365 | 1.144 | 18.284 | 8.354 | 0.071 | 4.106 | 814.809 |
| Mondego | 452.007 | 18.822 | 0.017 | 5.760 | 5.326 | 0.338 | 0.256 | 491.666 |
| Ouse | 10969.316 | 2.485 | 2.020 | 0.119 | 0.102 | 0.000 | 962.941 | 423.761 |
| Rolfsan | 55.807 | 0.020 | 25.972 | 1.134 | 1.624 | 0.000 | 1343.585 | 31.439 |
| Severn | 1003.194 | 14.447 | 0.308 | 0.078 | 0.671 | 0.004 | 126.309 | 608.963 |
| Tagus | 452.007 | 18.822 | 0.017 | 5.760 | 5.326 | 0.338 | 0.256 | 491.666 |
| Ula | 452.007 | 18.822 | 0.017 | 5.760 | 5.326 | 0.338 | 0.256 | 491.666 |

Table S1(cont.) –

| **Population** | **Mackerel** | **Grey mullet** | **Salmon** | **Twaite shad** | **Allis shad** | **Basking shark** | **Cod** | **Hake** |
| --- | --- | --- | --- | --- | --- | --- | --- | --- |
| Frome | 1003.194 | 14.447 | 0.308 | 0.078 | 0.671 | 0.004 | 126.309 | 608.963 |
| Gironde | 1085.738 | 39.365 | 1.144 | 18.284 | 8.354 | 0.071 | 4.106 | 814.809 |
| Mondego | 452.007 | 18.822 | 0.017 | 5.760 | 5.326 | 0.338 | 0.256 | 491.666 |
| Ouse | 10969.316 | 2.485 | 2.020 | 0.119 | 0.102 | 0.000 | 962.941 | 423.761 |
| Rolfsan | 55.807 | 0.020 | 25.972 | 1.134 | 1.624 | 0.000 | 1343.585 | 31.439 |
| Severn | 1003.194 | 14.447 | 0.308 | 0.078 | 0.671 | 0.004 | 126.309 | 608.963 |
| Tagus | 452.007 | 18.822 | 0.017 | 5.760 | 5.326 | 0.338 | 0.256 | 491.666 |
| Ulla | 452.007 | 18.822 | 0.017 | 5.760 | 5.326 | 0.338 | 0.256 | 491.666 |

|  | Frome-Uk | Mondego-Pt | Tagus-Pt | Ouse-Uk | Ulla-Spa | Rolfsan-Swe | Gironde-Fra | Severn-Uk |
| --- | --- | --- | --- | --- | --- | --- | --- | --- |
| Frome-Uk | * | 0.485 | 0.482 | **0.006** | 0.119 | **0.000** | 0.952 | 0.053 |
| Mondego-Pt | 0.012 | * | 0.999 | 0.99 | 0.999 | 0.999 | 0.999 | 0.999 |
| Tagus-Pt | 0.011 | 0.006 | * | 0.99 | 0.999 | 0.990 | 0.999 | 0.999 |
| Ouse-Uk | **0.010** | 0.007 | 0.006 | * | 0.999 | 0.248 | 0.999 | 0.998 |
| Ulla-Spa | 0.011 | 0.007 | 0.007 | 0.006 | * | 0.936 | 0.999 | 0.999 |
| Rolfsan-Swe | **0.016** | 0.011 | 0.0108 | 0.010 | 0.010 | * | 0.999 | 0.845 |
| Gironde-Fra | 0.012 | 0.007 | 0.008 | 0.007 | 0.007 | 0.010 | * | 0.999 |
| Severn-Uk | 0.013 | 0.008 | 0.008 | 0.008 | 0.008 | 0.010 | 0.008 | * |

Table S2 – Pairwise comparisons between sampling locations. F_ST_ values below diagonal and p-values above diagonal. Significant values are highlighted in bold.

| **Candidate locus** | **Access Number** | **% similarity** | **Alignment length** | **e-value** |
| --- | --- | --- | --- | --- |
| 1838 | AB275439.1 | 80.95 | 567 | 5.27E-116 |
|  | XM_032968544.1 | 88.00 | 300 | 4.22E-92 |
|  | KF318008.1 | 87.93 | 290 | 1.53E-86 |
|  | MN368861.1 | 87.71 | 236 | 1.21E-67 |
|  | AY577941.1 | 86.67 | 180 | 2.71E-44 |
| 3235 | XM_032965022.1 | 100.00 | 189 | 1.12E-90 |
| 13235 | XM_032979283.1 | 88.95 | 190 | 3.26E-56 |
|  | MN484596.1 | 83.87 | 248 | 4.22E-55 |
| 13283 | XM_032953481.1 | 100.00 | 222 | 5.05E-109 |
| 13513 | XM_032949755.1 | 83.84 | 229 | 5.50E-49 |
|  | AF464190.1 | 80.50 | 241 | 2.00E-38 |
| 14785 | XM_032949755.1 | 83.84 | 229 | 5.50E-49 |
|  | AF464190.1 | 80.50 | 241 | 2.00E-38 |
| 15374 | KF318011.1 | 88.65 | 467 | 6.28E-153 |
|  | AH007079.2 | 90.84 | 393 | 2.96E-141 |
|  | LC149809.1 | 88.19 | 364 | 3.07E-116 |
| 18413 | X92516.1 | 92.38 | 735 | 0 |
|  | LC149805.1 | 88.19 | 364 | 3.00E-116 |
| 23021 | XM_032944812.1 | 100.00 | 227 | 8.39E-112 |
|  | LS423642.1 | 92.75 | 207 | 1.14E-75 |
| 26013 | XM_032965631.1 | 96.12 | 309 | 2.61E-136 |
|  | XM_032965630.1 | 96.12 | 309 | 2.61E-136 |
| 27929 | X92516.1 | 91.97 | 735 | 0 |
|  | LC149805.1 | 87.09 | 364 | 1.17E-109 |

Table S3 – Accession numbers and blasting statistics for candidate loci
